# Surface Plasmon Resonance (SPR)-Based Workflow for High-Throughput Discovery of CD28-Targeted Small Molecules

**DOI:** 10.1101/2025.06.12.659248

**Authors:** Laura Calvo-Barreiro, Hossam Nada, Saurabh Upadhyay, Moustafa T. Gabr

## Abstract

CD28 is a critical costimulatory receptor involved in T cell activation and immune regulation, making it a compelling target for immunomodulatory therapies. Despite its therapeutic relevance, small molecule CD28 inhibitors remain largely underexplored. To address this gap, we developed a high-throughput screening (HTS) workflow using surface plasmon resonance (SPR) to identify novel CD28-targeted small molecules. To our knowledge, this work represents the first SPR-based HTS platform applied to the discovery of small molecules targeting a stimulatory immune checkpoint receptor. A chemical library composed of diverse 1,056 small molecules was screened using a 384-well format. Compounds were evaluated based on level of occupancy (LO), binding response, and dissociation kinetics, resulting in 12 primary hits (1.14% hit rate). Follow-up dose–response SPR screening confirmed micromolar-range affinities for three compounds. Molecular docking and 100 ns molecular dynamics (MD) simulations of the top hit, **DDS5**, revealed a stable complex with CD28, maintained by hydrogen bonding and a persistent interaction with Phe93. Functional validation using a competitive ELISA confirmed that **DDS5** inhibited the CD28–CD80 interaction. These results demonstrate that our SPR-based HTS platform is a robust and efficient strategy for discovering CD28-targeted small molecules. The integration of computational evaluation and orthogonal validation further underscores the potential of **DDS5** as an early-stage immunomodulatory agent.

---

T cell activation is a tightly regulated process that requires both antigen recognition and costimulatory signaling. Among the key costimulatory receptors, Cluster of Differentiation 28 (CD28) plays a central role in initiating and sustaining T cell responses^1^. In the absence of CD28 signaling, engagement of the T cell receptor (TCR) alone can result in T cell anergy, deletion, or regulatory differentiation—mechanisms essential for maintaining peripheral tolerance and preventing autoimmunity^1^. CD28 is constitutively expressed on naïve T cells and interacts with its classical ligands, CD80 (B7-1) and CD86 (B7-2), on antigen-presenting cells (APCs). This interaction amplifies TCR-mediated signaling by promoting the activation of downstream pathways such as PI3K–Akt, NF-κB, and MAPK, which collectively enhance interleukin-2 (IL-2) production, cell cycle progression, and anti-apoptotic responses^2^.

Dysregulation of CD28 signaling has been implicated in the pathogenesis of several immune-mediated diseases, including inflammatory bowel disease^3-5^ and rheumatoid arthritis^6^. Consequently, therapeutic strategies targeting CD28 have primarily focused on biologics, such as monoclonal antibodies and Fc fusion proteins, which block ligand binding and downstream signaling^7-10^. These approaches aim to dampen excessive T cell activation by preventing CD28 engagement with its ligands, thereby restoring immune balance in chronic inflammatory conditions. While these protein-based therapies have shown efficacy in certain contexts, they are associated with limitations including immunogenicity, limited tissue penetration, and complex manufacturing processes^11^. Moreover, the systemic nature of biologics can lead to off-target effects and prolonged immune suppression, necessitating more selective and controllable therapeutic alternatives^11^.

In contrast, small molecule inhibitors offer several advantages, including oral bioavailability, tunable pharmacokinetics, and reduced risk of anti-drug antibody formation^12^. Nevertheless, the development of CD28-targeted small molecule modulators has been hindered by the inherent structural and biophysical challenges of protein–protein interaction (PPI) interfaces, which are often characterized by shallow, flexible, and poorly defined binding topologies. These features limit the ability of small molecules to achieve high-affinity and selective binding. Recent advances in biophysical screening technologies have facilitated the discovery of small molecule modulators targeting PPIs. Among these, surface plasmon resonance (SPR) has emerged as a powerful, label-free technique that enables real-time detection of molecular interactions with high sensitivity and throughput. SPR is particularly well-suited for interrogating challenging targets such as immune checkpoint receptors, as it permits direct measurement of binding kinetics and affinities under near-physiological, solution-phase conditions. Moreover, its compatibility with fragment- and diversity-based libraries makes SPR an ideal platform for the early identification of novel chemotypes against traditionally undruggable targets.

To address the unmet need for small molecule modulators of CD28, we developed a high-throughput screening (HTS) platform based on SPR. Using this platform, we screened a diverse chemical library of small molecules and identified multiple hit compounds (1.14% hit rate) with micromolar binding affinities to CD28. The top candidate demonstrated functional inhibition of the CD28–CD80 interaction, supporting its potential as an immunomodulatory agent. To gain mechanistic insight into its mode of action, we performed molecular docking studies and molecular dynamics (MD) simulations, which revealed plausible binding poses at the CD28 interface that may sterically hinder ligand engagement.

We first defined and developed the affinity-based screening assay using SPR technology. The extracellular domain of the human CD28 protein (residues Asn19–Pro152, UniProt accession number P10747-1) was selected as the target ligand, as it represents the biologically active, glycosylated, disulfide-linked homodimeric form of the protein. This domain is responsible for ligand binding and is structurally well-suited for immobilization in SPR assays. As an essential first step, we optimized the immobilization strategy by selecting the most appropriate sensor chip format to ensure stable and reproducible ligand presentation for high-throughput interaction analysis.

Out of the diverse array of sensor chips compatible with the Biacore instrument, we selected the Sensor Chip CAP based on several key advantages. First, it enables the reversible capture of biotinylated molecules, facilitating chip regeneration and repeated use. Second, the Avitag™-labeled target protein exhibits exceptionally high binding affinity to the chip’s modified streptavidin (K_d_ = 4×10^−14^M, biotin-streptavidin^13^), ensuring stable immobilization with minimal dissociation over extended periods of time. Third, the Avitag™, a 15-amino acid peptide, is positioned downstream of a polyhistidine (His) tag at the C-terminus of the target protein. This configuration is not expected to interfere with the protein’s native structure or function, as both the His-tag and Avitag™ are relatively small and terminally located, thereby minimizing potential steric hindrance or conformational disruption.

We conducted a protein concentration scouting experiment ranging from 10 to 50 µg/mL and determined that 50 µg/mL was optimal (*data not shown*). This concentration enabled a ligand immobilization level (R_L_) of approximately 1,750 Response Units (RU), which in turn allowed us to achieve theoretical R_max_ values (maximum analyte-ligand interaction response) between 14 and 24 RU for our screened compounds (MW_L_ = 275-475 Da, respectively). Later, we optimized the assay buffer using a positive control, anti-CD28 antibody (reported IC_50_ ≈ 50 ng/mL in a cell-based assay^14^). We confirmed that 1x PBS-P+ (Cat # 28995084, Cytiva), with or without up to 2% DMSO, did not interfere with the expected protein-antibody binding affinity (*data not shown*). After verifying that DMSO concentrations up to 2% did not affect protein stability or function, we compared DMSO-supplemented 1x PBS-P+ with DMSO-supplemented 1x HBS-P+ (Cat# BR100671, Cytiva). No difference in RU was observed when a 2 µg/mL (saturating) dose of anti-CD28 antibody was tested (*data not shown*). Based on these findings, 50 µg/mL of His/Avitag™ human CD28 protein and a PBS-based buffer supplemented with 2% DMSO were selected for subsequent HTS of small molecules.

A 1,056-compound subset from the Discovery Diversity Set (DDS) library (Enamine) was selected to validate the SPR-based screening workflow. This library is particularly well-suited for targeting CD28, a transmembrane costimulatory receptor, due to its enrichment in chemotypes designed to engage G-Protein Coupled Receptors (GPCR)-like and protein–protein interaction (PPI)-like interfaces^15^. Such structural features are advantageous for binding to membrane proteins like CD28, which often present shallow, hydrophobic, or conformationally dynamic binding sites. The library’s emphasis on three-dimensional, sp^3^-rich scaffolds further enhances the probability of identifying ligands capable of accessing these challenging topologies. In addition, its diverse clustering strategy ensures broad coverage of chemical space, making it a robust starting point for both hit discovery and lead optimization in immunomodulatory drug development.

While a clean screen assay is often used to pre-filter compounds that exhibit nonspecific interactions with the sensor chip surface^16^, we elected to omit this step in our workflow. Compounds exhibiting such promiscuous behavior can be effectively identified during the single-concentration screen through elevated signals on the reference flow cell and atypical binding profiles. Moreover, nonspecific binding to the immobilized target may increase baseline responses but does not compromise target protein integrity or obscure the detection of true binders^17^. Given these considerations, and to streamline throughput, we proceeded directly with the primary screen, incorporating analytical flags to account for nonspecific and non-dissociating interactions during hit triage.

Subsequently, sourced from the 10 mM stock plates, we prepared the assay 384-well plates containing 100 µM of the small molecules in assay buffer supplemented with 2% DMSO. Additionally, negative control samples, containing only assay buffer with the same DMSO concentration but no small molecules, and positive control samples, containing anti-CD28 antibody at 2 µg/mL, were included in the 96-well plate where the remaining SPR reagents were also present. All plates were sealed with the appropriate foil to prevent sample evaporation throughout the duration of the experiment. Subsequently, the plates were placed into the Biacore hotel trays, and the HTS was carried out over a 19-hour period (*full details on the method in the Supporting Information*). After raw data collection, the following data analysis workflow was designed and applied to identify primary hit candidates for subsequent binding affinity experiments using the Biacore™ Insight Evaluation Software (Cytiva).

The first step involved applying the predefined evaluation method *LMW (Low Molecular Weight) screen using capture*, under the *Binding screen* tab, to the raw data obtained from the HTS. After applying solvent correction to all samples, we verified that the immobilization levels across the channels were consistent (ranging from 1,764.8 to 1,820.2 RU) and recorded the individual values for subsequent analysis. Under *Home > Properties > Run*, we entered the ligand information (Biotinylated human CD28 protein, MW 70,000 Da) along with the *capture levels* for each channel, as this information is required for further calculations. Within the *Evaluation - Binding* tab, we added all necessary parameters (as outlined in the formulas below) as independent columns in the data table. This setup enabled the use of the *Column Calculator* (within the *gear icon*) to apply the formulas needed for data analysis and hit selection. Specifically, R_max_ [Equation (1)] and Level of Occupancy [LO, Equation (2)].

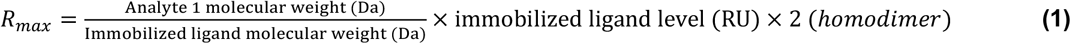

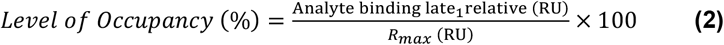

Once all necessary parameters were included for each screened compound, we proceed with data analysis. Under the *Home > Curve Markers* tab, we added markers to label each compound with one or several of the following categories: non-specific binder (NSB), non- or low-dissociating binder (NDB), high level of occupancy compound (High LO), and potential hit. Compounds that exhibited a relative response equal to or greater than 5 RU on the reference flow cell (*QC – Binding to Reference* tab) at the *Analyte Stability Early_1 Relative (RU)* time point were classified as NSB (123 out of 1,056 total screened compounds, 11.65% rate).

The typical behavior of small molecules is characterized by rapid association and dissociation with the target protein. Full association with the target ligand generally occurs within a few seconds, followed by a stable plateau phase as long as the small molecule continues to flow over the sensor chip. Once the flow stops, a rapid dissociation is expected. Therefore, compounds that exhibited minimal or no dissociation during the dissociation phase were flagged as NDB. For High LO classification, small molecules were divided into two groups: those with occupancy levels exceeding 200% were excluded from further consideration (1 out of 1,056 compounds, 0.09% rate), while those with occupancy between 100% and 200% were flagged. Compounds with occupancy levels between 50% and 100% were flagged as potential hits.

Taking all these criteria into account, those small molecules that had no NSB flag and their LO was between 50% and 200% were selected for subsequent binding affinity experiments (12 out of 1,056 compounds, 1.14% hit rate) (Table 1, Figure 1). Among these, nine compounds presented a LO between 50% and 100% and no additional flags (Table 1, Figure 1: solid lines). Compound **DDS1** exhibited a higher-than-expected LO (189.70%) and a slow-dissociating behavior (Table 1, Figure 1: dashed line). Compounds **DDS7** and **DDS10** fell within the acceptable LO but displayed a slow-dissociating behavior (Table 1, Figure 1: dotted lines). This HTS and subsequent data analysis workflow enabled the rapid classification, ranking, and prioritization of primary hits for follow-up binding affinity experiments. Selection was based on each compound’s binding response and behavior toward CD28, while excluding small molecules exhibiting aberrant or undesirable binding characteristics.

**Table 1.** List of primary small molecule hits identified through high-throughput screening. Compounds were selected for follow-up binding affinity studies if they exhibited a Level of Occupancy (%) greater than 50% and less than 200% and were not flagged as non-specific binders. For each compound, the table includes molecular weight (MW), analyte binding level (corresponding to the *Analyte Binding Rate_1 Relative* [RU] at the end of the association phase), ligand name and MW, immobilized ligand level, the calculated R_max_ value, and the corresponding Level of Occupancy

| Analyte | Analyte MW (Da) | Analyte Binding Level (RU) | Ligand | Ligand MW (Da) | Immobilized Ligand Level (RU) | Rmax (RU) | Level of Occupancy (%) |
| --- | --- | --- | --- | --- | --- | --- | --- |
| DDS1 | 341.33 | 33.2 | Biotinylated Human CD28 Protein | 70,000 | 1792.5 | 17.48 | 189.70 |
| DDS2 | 298.26 | 14.4 |  |  | 1776.8 | 15.14 | 95.08 |
| DDS3 | 288.30 | 13.8 |  |  | 1786.5 | 14.72 | 93.47 |
| DDS4 | 321.39 | 12.2 |  |  | 1792.5 | 16.46 | 74.27 |
| DDS5 | 305.42 | 11.3 |  |  | 1798.1 | 15.69 | 72.12 |
| DDS6 | 305.37 | 10.7 |  |  | 1798.1 | 15.69 | 67.96 |
| DDS7 | 301.30 | 9.6 |  |  | 1786.5 | 15.38 | 62.35 |
| DDS8 | 444.52 | 12.3 |  |  | 1780.8 | 22.62 | 54.38 |
| DDS9 | 335.33 | 9.0 |  |  | 1764.8 | 16.91 | 53.51 |
| DDS10 | 300.42 | 8.0 |  |  | 1776.8 | 15.25 | 52.46 |
| DDS11 | 309.38 | 8.3 |  |  | 1792.8 | 15.85 | 52.16 |
| DDS12 | 364.44 | 9.5 |  |  | 1792.5 | 18.66 | 50.77 |

**Figure 1.**
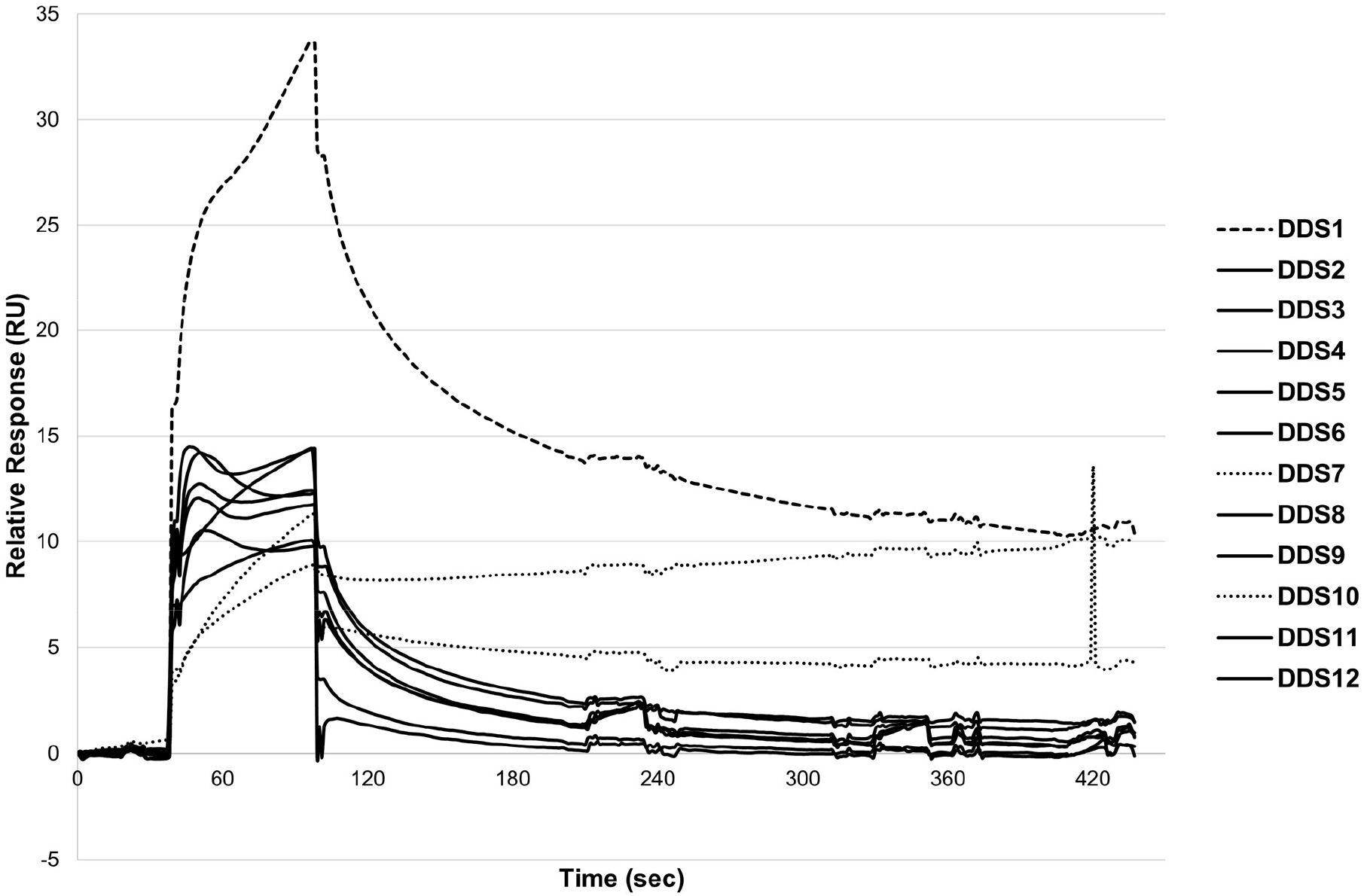
Sensorgrams of primary small molecule hits identified through HTS and selected for follow-up binding affinity studies. Sensorgrams represent the analysis step within the designed HTS method used to evaluate 1,056 compounds. All compounds were injected over the sensor chip at 100 µM with a 60-second association phase and a 100-second dissociation phase at a flow rate of 30 µL/min in running buffer (1x PBS-P+ supplemented with 2% DMSO). A wash step with 50% DMSO in Milli-Q water and a carry-over control using running buffer (20 seconds at 30 µL/min) followed, both bypassing the sensor chip flow cells. Compounds were selected for follow-up binding affinity studies if they exhibited a Level of Occupancy (%) greater than 50% and less than 200% and were not flagged as non-specific binders. Sensorgrams are color-coded as follows: solid black lines represent compounds with LO between 50% and 100% and no additional flags; dashed black lines indicate compounds with LO between 100% and 200% and slow-dissociating behavior; and dotted black lines represent compounds with LO between 50% and 100% but exhibiting slow-dissociating behavior.

To further characterize the binding interactions of the identified CD28 binders, we investigated whether each compound exhibited a dose-dependent response. Using SPR, we successfully measured K_d_ values in the micromolar range for three compounds (Table 2, Figure 2). Additionally, two compounds demonstrated weaker binding affinities, with K_d_ values in the low millimolar range (*data not shown*). The remaining compounds did not display a clear dose-response relationship (Table 2).

**Table 2.**
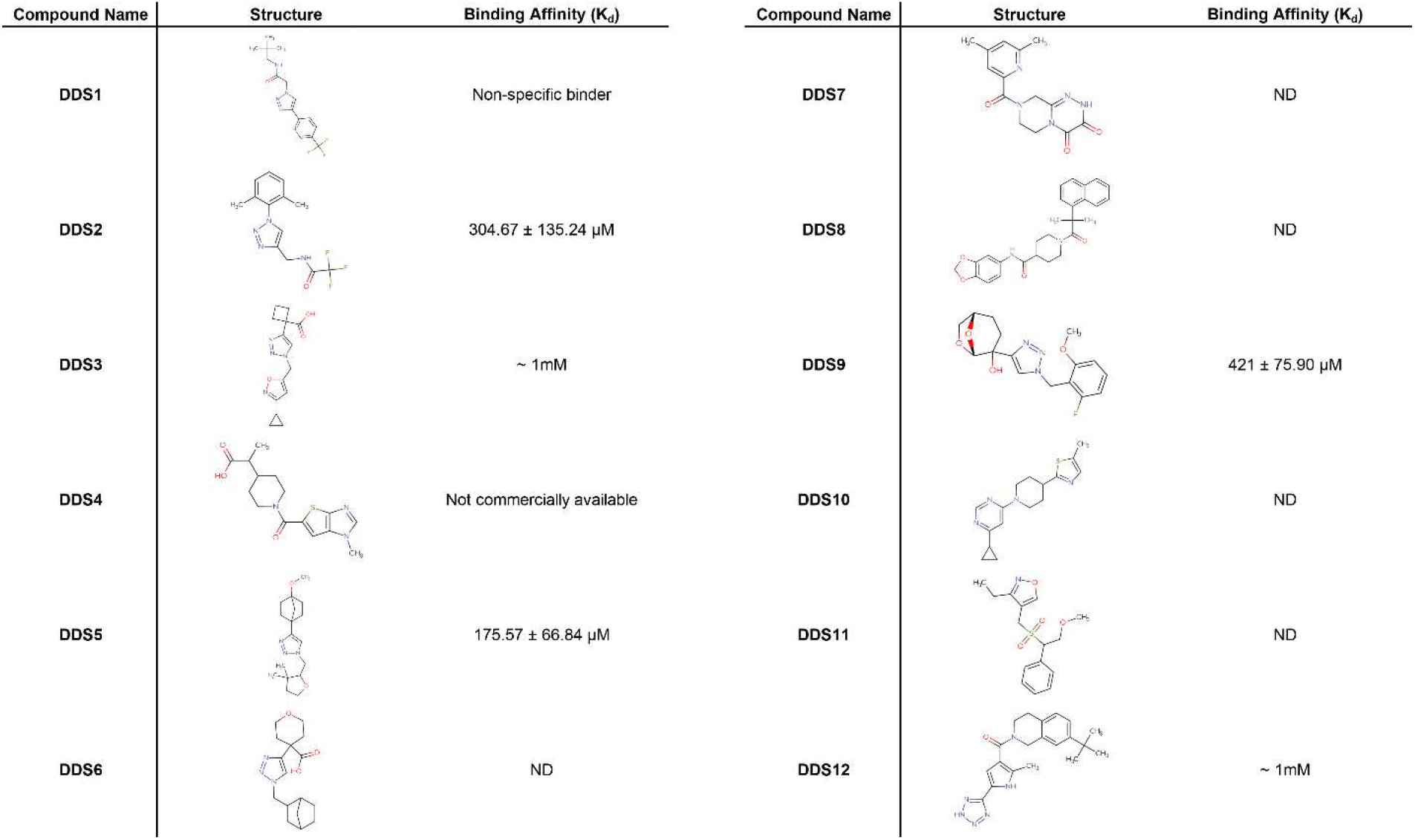
Binding affinities and structural information of selected primary small-molecule hits. This table summarizes the binding affinities (K_d_) and chemical structures of all compounds tested for interaction with the CD28 protein. Binding affinities are reported in micromolar (µM) or millimolar (mM) concentrations, with standard deviations where available. *ND* indicates that the binding affinity was not determined due to lack of dose-response effect. *Non-specific binder* denotes compounds that exhibited a dose-response effect but did not show selective binding to the active flow cell.

| Compound Name | Structure | Binding Affinity ($K_d$) | Compound Name | Structure | Binding Affinity ($K_d$) |
| --- | --- | --- | --- | --- | --- |
| DDS1 |  | Non-specific binder | DDS7 |  | ND |
| DDS2 | | $304.67 \pm 135.24 \mu\text{M}$ | DDS8 | | ND |
| DDS3 | | $\sim 1\text{mM}$ | DDS9 | | $421 \pm 75.90 \mu\text{M}$ |
| DDS4 |  | Not commercially available | DDS10 |  | ND |
| DDS5 | | $175.57 \pm 66.84 \mu\text{M}$ | DDS11 | | ND |
| DDS6 | | ND | DDS12 | | $\sim 1\text{mM}$ |

**Figure 2.**
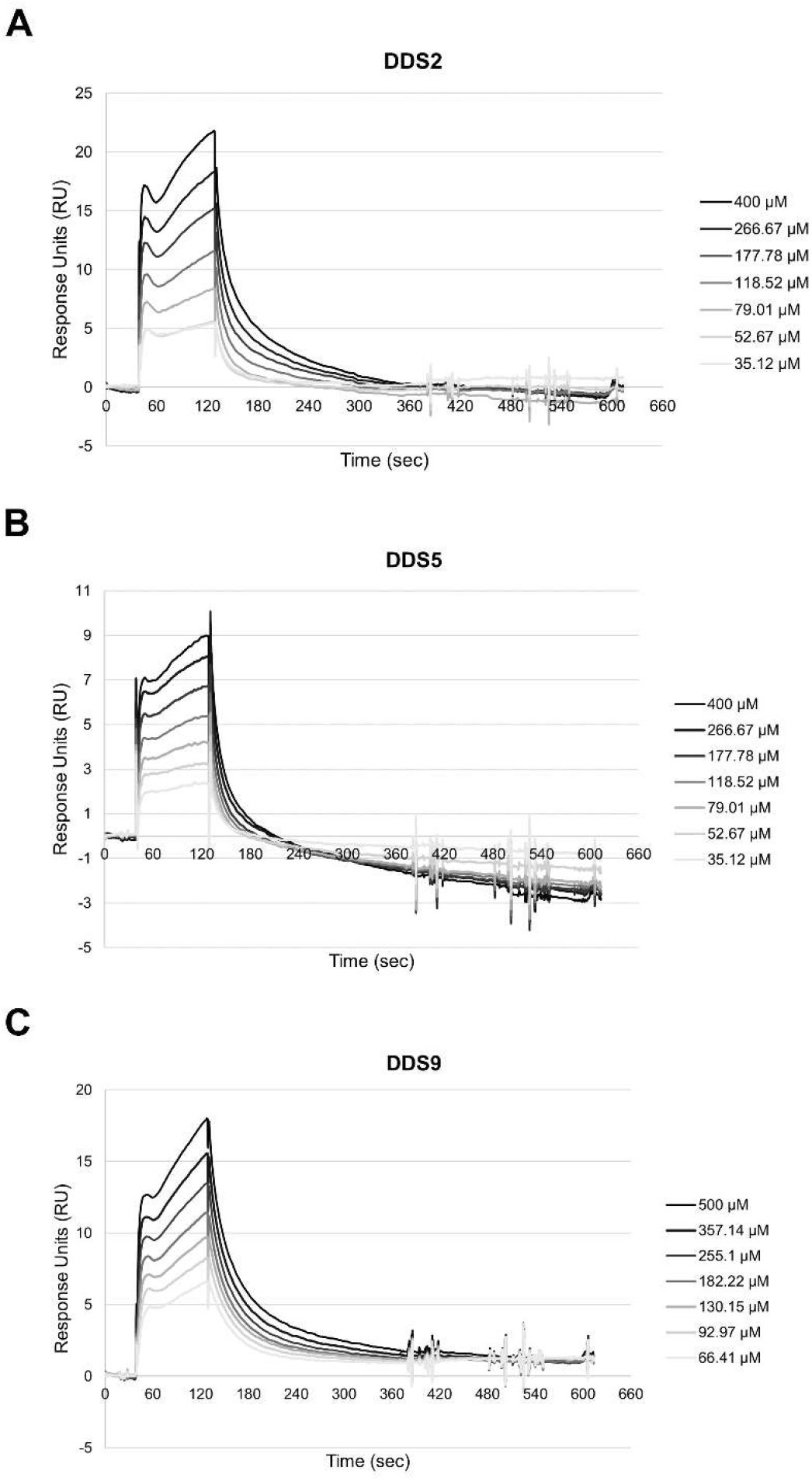
Binding affinity of selected small molecules to immobilized human CD28 protein. A. Serial dilutions of compound **DDS2** (ranging from 400 to 35.12 µM, including a 0 µM reference; 1.5-fold dilution series) were injected onto a Sensor Chip CAP with immobilized human CD28 protein using a multi-cycle kinetics method. **B**. Serial dilutions of compound **DDS5** (ranging from 400 to 35.12 µM, including a 0 µM reference; 1.5-fold dilution series) were similarly injected using the same method. **C**. Serial dilutions of compound **DDS9** (ranging from 500 to 66.41 µM, including a 0 µM reference; 1.4-fold dilution series) were also injected under the same conditions. The sensorgrams show relative Response Units (RU) over time during a 90-second association phase and a 240-second dissociation phase, representing one of three independent experiments. Binding curves were analyzed using non-linear curve fitting based on steady-state affinity analysis.

Following the identification of three CD28 binders with defined binding affinities, we sought to characterize their binding mode to CD28 and evaluate their potential to inhibit the CD28–CD80 interaction. As a proof of concept to validate the robustness of our HTS platform, the top-ranking compound, **DDS5**, was selected for further mechanistic and functional validation experiments.

The co-stimulatory receptor CD28 and its inhibitory counterpart CTLA-4 share a common ligand binding mechanism through their interaction with B7 family proteins (CD80/CD86)^1^. Previous crystallographic studies have established that the primary ligand binding site for both CD28 and CTLA-4 is located within a conserved region comprising residues 99-104^18^. Mutagenesis experiments have demonstrated that alterations within this sequence result in a substantial reduction (>90%) in ligand binding affinity, confirming its critical role in receptor-ligand recognition^18^. However, the primary ligand binding region of CD28 presents significant challenges for small molecule drug development. Analysis of the three-dimensional structure (PDB ID: 1YJD)^18^ revealed that the CD28-CD80 binding interface is characterized by a relatively shallow, extended surface lacking the well-defined binding clefts typically required for high-affinity small molecule interactions. Such flat protein-protein interaction surfaces are notoriously difficult to target with drug-like compounds due to their limited capacity to provide the multiple contact points necessary for selective binding.

Although the primary ligand-binding site of CD28 presents significant challenges for small-molecule intervention due to its shallow and extended topology, structural analysis has identified a secondary binding pocket with more favorable features for small molecule targeting. This CD28-specific cavity is positioned adjacent to the canonical ligand-binding region and is formed by a discontinuity in β-strand G. The pocket is delineated by key residues including His38, Phe93, Lys95, Asp106, and Lys109 forming the surrounding walls, with Val5, Asn107, and Ser110 comprising the base of the binding site (Figure 3A, highlighted region). These structural attributes suggest that the pocket may serve as a viable target for structure-based design of small molecule CD28 modulators.

**Figure 3.**
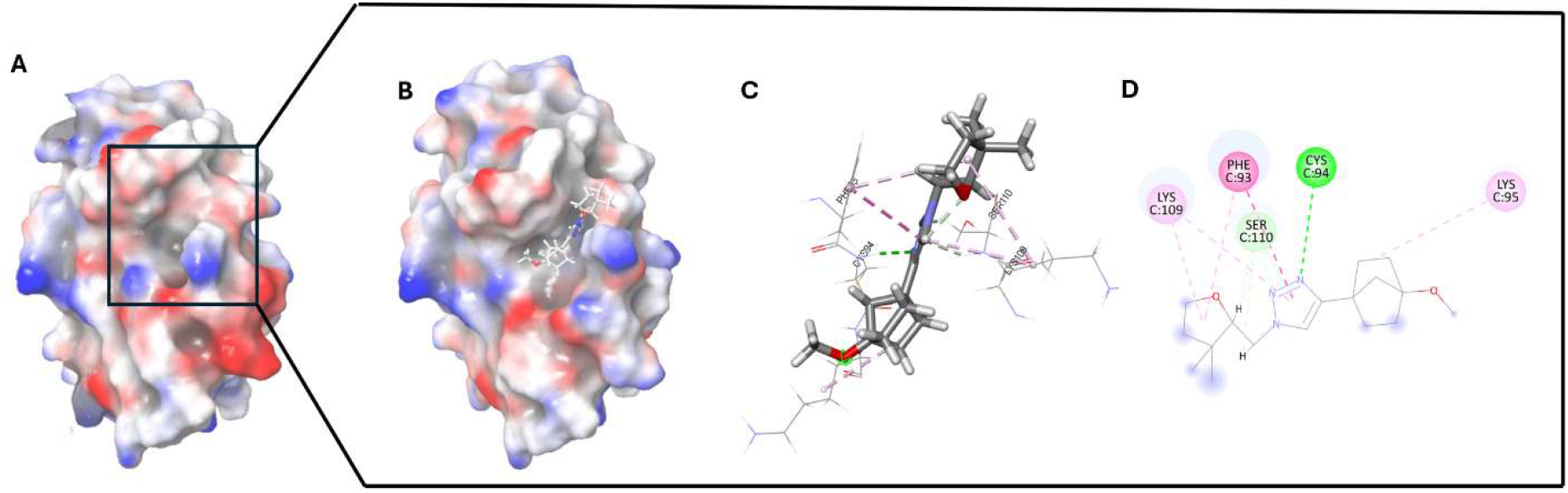
Structural analysis and molecular docking of CD28-DDS5 binding site. A. Crystal structure of CD28 (PDB ID: 1YJD) showing the primary ligand binding site and the identified secondary binding pocket (enclosed in a square). The secondary pocket is formed by discontinuity in β-strand G and surrounded by key residues His38, Phe93, Lys95, Asp106, and Lys109. **B**. Docked conformation of compound **DDS5** within the identified binding pocket, showing optimal fit and induced conformational changes. **C**. 3D bound complex of **DDS5** with CD28. **D**. Detailed view of **DDS5** binding interactions, highlighting the critical hydrogen bond formation with Cys94 and multiple hydrophobic contacts with surrounding residues.

The therapeutic relevance of targeting regions adjacent to primary binding sites is supported by precedent from CTLA-4 research. Structural studies of the anti-CTLA-4 antibody tremelimumab have demonstrated that nine of the ten residues within the heavy chain complementarity-determining region 3 (HCDR3) loop (residues 101-110) are directly involved in CTLA-4 binding^19^. These findings led to the hypothesis that cyclic peptides derived from the HCDR3 sequence could effectively block CD80/CD86 binding to CTLA-4^19^. Given the high degree of sequence and structural homology between CTLA-4 and CD28, it is plausible that analogous binding regions on CD28 may similarly be exploited for therapeutic modulation. In particular, the identified secondary pocket on CD28—composed of residues His38, Phe93, Lys95, Asp106, Asn107, Lys109, and Ser110—may represent a viable target for small-molecule intervention.

To further evaluate this hypothesis, we employed the SiteMap module within the Maestro software suite (Schrödinger). SiteMap utilizes physics-based scoring criteria to identify and rank potential binding pockets while filtering out regions unlikely to exhibit druggable properties^20^. Consistent with our manual structural analysis, SiteMap ranked the secondary pocket adjacent to the canonical ligand-binding site as the second most favorable site for small-molecule binding, reinforcing its potential as a novel druggable interface on CD28 (*data not shown*).

The top hit compound, **DDS5**, which exhibited the strongest binding affinity to CD28, was subjected to molecular docking analysis using the Induced Fit Docking (IFD) protocol. This approach accounts for the flexibility of both the ligand and the receptor, allowing for realistic conformational adjustments during the binding process^21^. **DDS5** demonstrated favorable binding characteristics within the previously identified secondary pocket (Figure 3B). The docking results suggested that the compound induces local conformational rearrangements within the binding site, optimizing the geometry to accommodate the ligand. Visual inspection of the predicted binding pose revealed the formation of a key hydrogen bond with Cys94, along with multiple hydrophobic interactions involving adjacent residues (Figure 3D). The combination of polar and nonpolar contacts contributes to a well-anchored and specific binding mode, supporting the potential of **DDS5** as a structurally viable modulator of CD28 function.

To evaluate the stability of the predicted protein–ligand complex, we performed two independent 100 ns MD simulations: one for unbound CD28 and the other for CD28 in complex with **DDS5**. Throughout the simulation period, **DDS5** remained stably bound to CD28, as evidenced by the root mean square deviation (RMSD) trajectory (Figure 4A, blue line). RMSD analysis revealed an average deviation of approximately 1.3 Å, indicating minimal structural perturbation and a highly stable binding conformation. To further support the stability of the interaction, protein–ligand contact analysis was conducted, which demonstrated that **DDS5** maintained persistent interactions with Phe93 across the full 100 ns simulation (Figure 4B). This sustained contact highlights both the structural integrity of the complex and the critical role of Phe93 as a potential anchor residue for ligand recognition and stabilization.

**Figure 4.**
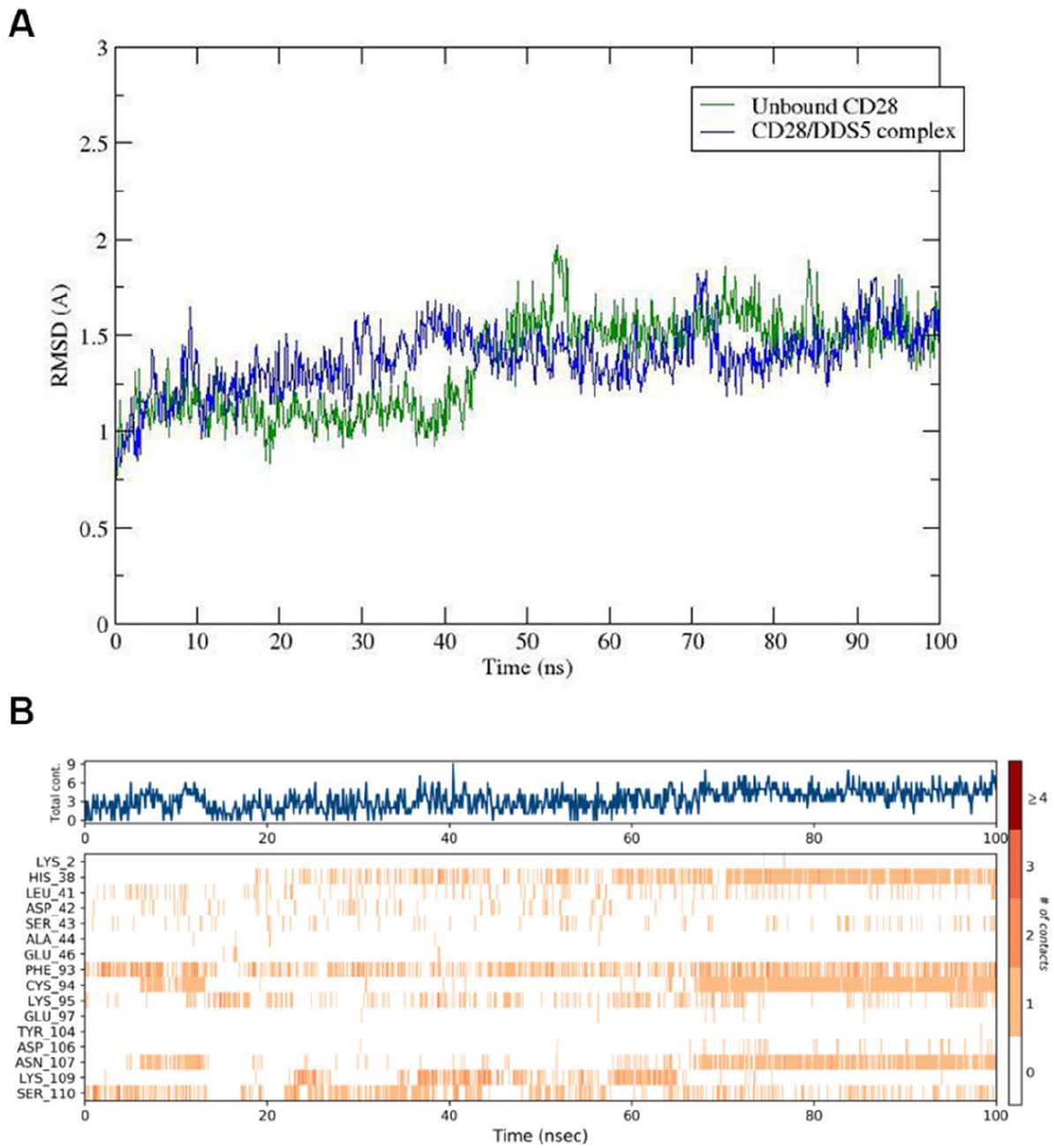
MD simulation analysis of compound DDS5 stability in CD28 binding site. A. Root mean square deviation (RMSD) trajectory of **DDS5**-CD28 complex over 100 ns simulation (blue line), demonstrating stable binding with average RMSD of ∼1.3 Å. **B**. Protein-ligand contact histogram for **DDS5** showing sustained interaction with Phe93 throughout the simulation period, confirming binding stability and the critical role of this residue.

Finally, we performed a quantitative assessment of the CD28-CD80 disruption by our **DDS5** candidate using an ELISA-based approach. As previously discussed, disrupting the CD28-CD80 interaction offers a promising strategy to modulate T cell co-stimulation, with therapeutic implications in not only in autoimmunity, but also in transplant rejection, and cancer^22-25^. In pathological conditions involving hyperactive immune responses, blocking this pathway can help suppress excessive inflammation and restore immune balance. To evaluate the ability of small molecules to interfere with this interaction, we conducted a competitive ELISA using immobilized CD28 and biotinylated CD80 as the ligand. Compound **DDS5** was tested across a range of concentrations and exhibited a dose-dependent inhibition of CD80 binding with an IC_50_ value of 332 µM (Figure 5). This finding is consistent with its moderate binding affinity (K_d_ = 175.57 ± 66.84 µM). Together, these results provide confirmation that our top compound directly engages CD28 and can interfere its binding with CD80, supporting its potential as an early-stage inhibitor for further optimization.

**Figure 5.**
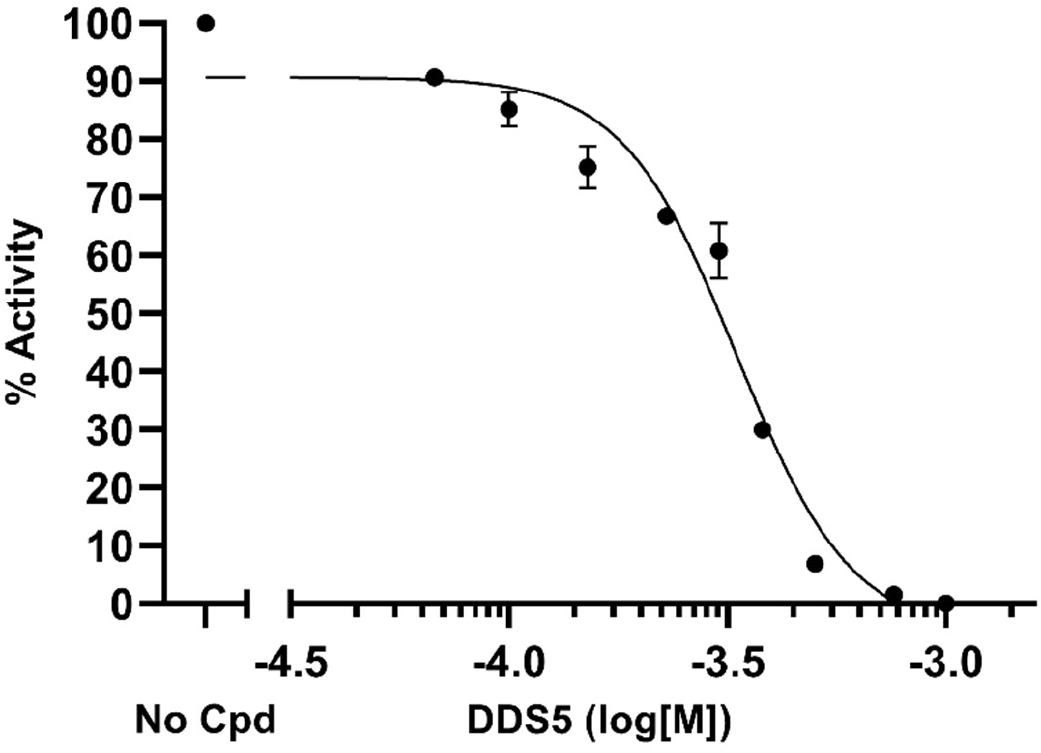
Inhibition of CD28-CD80 interaction by compound DDS5. Dose-dependent inhibition of CD28-CD80 binding by compound **DDS5** was measured using a competitive ELISA assay. The compound demonstrated an IC_50_ value of 332 µM. The maximum luminescence signal with no compound added was considered as 100% activity, while the minimum luminescence observed at the highest compound concentration was considered no activity. All intermediate values were normalized accordingly between these two reference points. IC50 calculations were performed using GraphPad Prism, applying the *log(inhibitor) vs. response – Variable slope (four-parameter)* nonlinear regression model. The graph shows the raw data from a single experiment with two technical replicates. The data are presented as the mean ± standard deviation.

In this study, we report the first successful implementation of a SPR-based HTS platform for the identification of small-molecule modulators targeting CD28—a historically challenging immune checkpoint receptor. By leveraging recent technological advancements in SPR, including enhanced sensitivity, throughput, and data resolution, we established a robust, label-free, and solution-phase method capable of directly quantifying binding kinetics and affinities. This makes SPR particularly advantageous for interrogating challenging targets such as CD28, which present flat, dynamic, and poorly defined PPI interfaces.

Our platform enabled the rapid and reproducible screening of a structurally diverse library of 1,056 compounds, culminating in a hit rate of 1.14%, with 5 of 12 primary hits (41.7%) exhibiting dose-dependent binding and quantifiable affinities. Notably, the single-concentration HTS campaign was completed within a single day, underscoring the operational efficiency and reliability of the workflow. The integration of biophysical screening with computational modeling and functional validation represents a comprehensive strategy for triaging hits early in the discovery process.

Our top hit compound, **DDS5**, demonstrated micromolar binding affinity, formed a structurally stable complex with CD28 as shown by MD simulations, and inhibited CD28–CD80 binding in a competitive ELISA. These results provide the first functional evidence of CD28–CD80 disruption by a non-peptidic small molecule. Docking studies further suggested a mechanism involving steric hindrance within a previously unexploited secondary pocket on CD28, providing a rational foundation for future structure–activity relationship (SAR) and hit-to-lead optimization campaigns.

Beyond the specific case of CD28, this study underscores the broader utility of SPR as a central platform in modern drug discovery. SPR not only accelerates early-stage screening and target validation but also enables detailed mechanistic interrogation, lead prioritization, and risk reduction in downstream development. Compared to other biophysical techniques such as thermal shift assays, X-ray crystallography, and NMR, SPR offers unique advantages—real-time kinetic resolution, low sample consumption, and broad molecular compatibility—making it indispensable for fragment-based design, mode-of-action studies, and selectivity profiling.

Taken together, our work establishes a blueprint for SPR-enabled discovery of small-molecule immune modulators and highlights its potential to expand the therapeutic landscape for T cell–targeted immunotherapies. Furthermore, the approach may be extended to other costimulatory pathways, such as ICOS, and integrated into multi-target discovery pipelines aimed at addressing complex immune-driven diseases. By demonstrating the druggability of CD28 with small molecules, this work contributes to redefining what is possible in the chemical modulation of immune checkpoint biology.

## ASSOCIATED CONTENT

Supporting Information

The Supporting Information is available free of charge on the ACS Publications website.

Experimental procedures and chromatographic and mass spectra data for the identified hit compounds (PDF)

## Author Contributions

The manuscript was written through contributions of all authors. All authors have given approval to the final version of the manuscript.

## AUTHOR INFORMATION

Corresponding Author

*

## ABBREVIATIONS

CAP: capture sensor chip
CD28: cluster of differentiation 28
CD80: cluster of differentiation 80 (B7-1)
CD86: cluster of differentiation 86 (B7-2)
DDS: Discovery Diversity Set
ELISA: enzyme-linked immunosorbent assay
GPCR: G protein-coupled receptor
HTS: high-throughput screening
IC_50_: half-maximal inhibitory concentration
Kd: equilibrium dissociation constant
LO: level of occupancy
MD: molecular dynamics
PPI: protein–protein interaction
Rmax: maximum theoretical response
RU: response units
SPR: surface plasmon resonance.

## Notes

The authors declare no competing financial interests.

## Acknowledgments

This work was supported by the National Institute of Diabetes and Digestive and Kidney Diseases (NIDDK) under grant number R01DK137299. The author gratefully acknowledges Michael B. Murphy (Cytiva) for his expert guidance and consistent support in interpreting complex sensorgram data obtained using the Biacore™ 8K instrument. His recommendations on methodological adjustments were instrumental in optimizing the experimental design and data quality throughout this study. Finally, we would like to thank the Fisher Drug Discovery Resource Center of Rockefeller University (RRID:SCR_020985) for providing access to the Cytiva Biacore 8K instruments.

## Table of Contents graphic

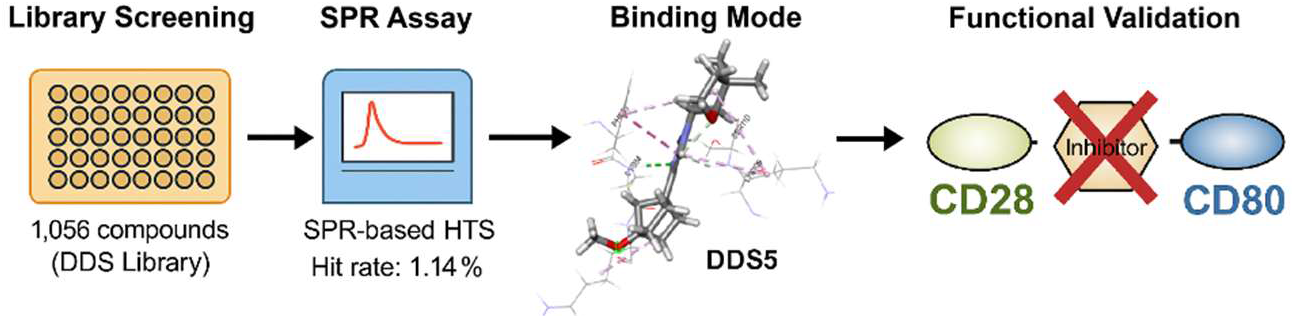

## Notes

### Competing Interest Statement

The authors have declared no competing interest.

