## Supporting Information for "Surface Plasmon Resonance (SPR)-Based Workflow for High-Throughput Discovery of CD28-Targeted Small Molecules"

### 1. Experimental Methods

#### 1.1. Screened Library

A subset of 1,056 compounds held within the Discovery Diversity Set Library (Cat# DDS-10-10-Y-10, Enamine, Kyiv, Ukraine) was chosen as suitable candidate library for the validation of the designed workflow. All screened compounds were contained in 384-well plates and dissolved in 100% DMSO at a concentration equal to 10 mM. Libraries were stored at -80°C upon arrival and until use.

After High-Throughput Screening (HTS) of stock plates, new lyophilized material for every selected small molecule candidate was acquired from Enamine, reconstituted in DMSO at a 50 mM stock concentration and stored at -30°C until use.

#### 1.2. High-Throughput Screening Using SPR

A single-dose (100 µM) HTS of a focused library comprising 1,056 compounds was conducted using a Biacore™ 8K instrument (Cytiva, Marlborough, MA, USA) at 25 °C. The running buffer consisted of 1x PBS-P+ (Cat# 28995084, Cytiva) supplemented with 2% DMSO. A custom method was developed in Biacore^TM^ 8K Control Software (Cytiva) to perform a binding-level screening assay, consisting of the following steps:

The procedure began with an *Initial Regeneration*, which involved three regeneration cycles (60 seconds each at 10 µL/min) using a solution prepared by combining 3 parts of Regeneration Stock 1 and one part of Regeneration Stock 2 from the Biotin CAPture Kit, Series S (Cat# 28920234, Cytiva). This solution effectively removes any Biotin CAPture Reagent, immobilized ligand, and bound analyte from the chip surface, enabling chip reuse. A subsequent wash with running buffer was applied through the flow system, bypassing the sensor chip flow cells.

During *System Startup*, Biotin CAPture Reagent (from the Biotin CAPture Kit, Series S) was diluted 1:1 with 1x HBS-EP+ buffer (Cat# BR100669, Cytiva) and injected for 300 seconds at 2 µL/min without a dissociation phase. This was followed by a capture cycle in flow path 2 (active flow cell) using 1x PBS-P buffer (300 seconds at 10 µL/min), and a flow system wash with 50% isopropanol and 50 mM NaOH/1 M NaCl, excluding the sensor chip flow cells. Analyte binding was assessed using 1x PBS-P with 2% DMSO (null analyte), with a 60-second contact time and 100-second dissociation at 30 µL/min. Regeneration was then performed for 120 seconds at 10 µL/min, followed by a wash with running buffer.

For *CD28 Immobilization*, Biotin CAPture Reagent was again diluted 1:1 with 1x HBS-EP+ and injected over both flow cells (300 seconds, 2 µL/min, no dissociation). Biotinylated human CD28 protein (His, Avitag™, Cat# CD8-H82E5, Acro Biosystems, Newark, DE, USA) was captured in flow path 2 (300 seconds, 10 µL/min, 50 µg/mL), achieving an immobilization level of approximately 1,750 RU. Before being added to the corresponding plate, target protein was centrifuged at 15,000 rpm for 10 minutes and 4°C to remove any protein aggregates. This was followed by a flow system wash with 50% isopropanol and 50 mM NaOH/1 M NaCl, and five additional washes with running buffer, excluding the sensor chip flow cells.

The *Small Molecule Binding Assay* involved injecting candidate small molecules at 100 µM with a 60-second contact time and 100-second dissociation at 30 µL/min. A wash with 50% DMSO in Milli-Q water and a carry-over control using running buffer (20 seconds at 30 µL/min) followed, both excluding the sensor chip flow cells. A negative control (1x PBS-P with 2% DMSO) was included every 16 cycles, followed by the same wash and carry-over steps. Compounds were screened using 384-well plates (Cat# 781280, Greiner Bio-One, Kremsmünster, Austria), which were centrifuged at 1,000 rpm for 1 minute at room temperature and sealed with the recommended film (Cat# BR100577, Cytiva) prior to loading into the instrument.

*Solvent Correction* and *Positive Control* steps were performed to ensure data accuracy and system validation. Solvent corrections (four solutions per cycle) were conducted before and after the positive control to account for DMSO-related refractive index changes. The positive control involved injecting an anti-CD28 antibody (Cat# K123A, Promega, Madison, WI, USA) at 2 µg/mL with a 180-second contact time and 600-second dissociation at 30 µL/min.

To conclude the assay, a *Final Regeneration* was performed using a 120-second regeneration step at 10 µL/min, followed by three washes with running buffer, again excluding the sensor chip flow cells.

*Data Acquisition* were carried out using Biacore™ 8K Control Software (Cytiva), with subsequent *analysis* performed using a custom workflow (*detailed in the Results section*) in Biacore™ Insight Evaluation Software (Cytiva). This workflow enabled the identification, ranking, and selection of primary hits based on their binding responses to CD28, while filtering out compounds with aberrant or undesirable binding characteristics.

#### 1.3. Binding Affinity Screening Using SPR

A multi-cycle kinetic assay was performed using a Biacore™ 8K instrument (Cytiva) at 25 °C to characterize the binding kinetics of primary hits identified during the previously performed single-dose HTS. The running buffer consisted of 1x PBS-P+ (Cat# 28995084, Cytiva) supplemented with 2% DMSO. A custom method was developed in Biacore™ 8K Control Software (Cytiva) to execute the assay, comprising the following steps:

An *Initial Regeneration* was performed as described in the previous methods section, using three regeneration cycles (60 seconds each at 10 µL/min) with a solution prepared from the Biotin CAPture Kit, Series S (Cat# 28920234, Cytiva). This step, which effectively removes residual Biotin CAPture Reagent, immobilized ligand, and bound analyte to enable chip reuse, was maintained without modification from the previously detailed protocol. A subsequent wash with running buffer was applied through the flow system, bypassing the sensor chip flow cells.

*System Startup* began with the injection of Biotin CAPture Reagent (from the Biotin CAPture Kit, Series S), diluted 1:1 with 1x HBS-EP+ buffer (Cat# BR100669, Cytiva), for 300 seconds at 2 µL/min without a dissociation phase. A capture cycle was then executed in flow path 2 using 1x PBS-P buffer (300 seconds at 10 µL/min). The system was subsequently washed with 50% isopropanol and 50 mM NaOH/1 M NaCl, bypassing the sensor chip flow cells. Analyte binding was assessed using 1x PBS-P with 2% DMSO (null analyte), with a 60-second contact time and 100-second dissociation at 30 µL/min. A regeneration step followed (120 seconds at 10 µL/min), and the system was washed again with running buffer.

*CD28 Immobilization* was carried out by injecting diluted Biotin CAPture Reagent over both flow cells (300 seconds, 2 µL/min, no dissociation). Biotinylated human CD28 protein (His, Avitag™, Cat# CD8-H82E5, Acro Biosystems) was captured in flow path 2 (300 seconds, 10 µL/min, 50 µg/mL), achieving an immobilization level of approximately 1,750 RU. This was followed by a wash with 50% isopropanol and 50 mM NaOH/1 M NaCl, excluding the sensor chip flow cells.

*Pre-Analysis Washes* were performed to stabilize the baseline. Five cycles of 1x PBS-P with 2% DMSO were injected (60-second contact, 120-second dissociation at 30 µL/min) over both flow cells, followed by five washes with running buffer, bypassing the sensor chip flow cells.

*Multi-Cycle Kinetic* was conducted to determine the binding kinetics of the small molecule hits. One buffer blank and seven ascending concentrations of each analyte (ranging from 35.12 µM up to 500 µM) were injected sequentially, with each cycle consisting of a 90-second contact time and a 240-second dissociation phase at 30 µL/min. Buffer blank responses were later subtracted from analyte curves to correct for baseline drift. Between cycles, the flow system, except the sensor chip flow cells, was washed with 50% DMSO in Milli-Q water, followed by a carry-over control injection (20 seconds at 30 µL/min, running buffer only). Compounds were screened using 96-well plates (Cat# 650201, Greiner Bio-One), which were centrifuged at 1,000 rpm for 1 minute at room temperature and sealed with the recommended film (Cat# 28975816, Cytiva) prior to loading into the instrument.

To maintain data integrity and validate system performance, *Solvent Correction* and *Positive Control* steps were systematically incorporated. Solvent Correction was carried out using four DMSO-based solutions per cycle (ranging 1.5% up to 3%), both preceding and following the positive control, to compensate for refractive index variations associated with DMSO. For the Positive Control, an anti-CD28 antibody (Cat# K123A, Promega) was injected at a concentration of 2 µg/mL. The injection was performed at a flow rate of 30 µL/min, with a 180-second association phase followed by a 600-second dissociation phase.

At the conclusion of the assay, a *Final Regeneration* step was applied. This involved a 120-second regeneration cycle at 10 µL/min, succeeded by three sequential washes with running buffer.

*Data were acquired* using Biacore™ 8K Control Software (Cytiva) and analyzed by non-linear curve fitting using a steady-state affinity analysis in Biacore™ Insight Evaluation Software (Cytiva).

#### 1.4. Binding Site Characterization and Ligand Docking

The three-dimensional structure of human CD28 in complex with the Fab fragment of a mitogenic antibody was retrieved from the Protein Data Bank (PDB ID: 1YJD). The downloaded structure underwent comprehensive energy minimization and structural refinement using the Maestro Schrödinger Suite employing the OPLS3e force field. Putative binding hotspot residues within the CD28 interface were systematically identified using the SiteMap wizard integrated within Maestro. The selected hit compound was docked into the computationally identified binding site using the Induced Fit Docking (IFD) protocol implemented in Maestro Schrödinger Suite (version 2021.2). The IFD approach was selected to accommodate the inherent flexibility of both ligand and protein binding site, enabling realistic conformational adjustments during the docking procedure. Ligand structure was prepared using the LigPrep module within Maestro, while protein structure preparation was performed using the Protein Preparation Wizard. The IFD protocol consisted of three sequential steps: an initial Glide docking phase, followed by Prime-based refinement of the protein-ligand complex, and concluded with a final Glide redocking step to optimize the final binding pose. Molecular dynamics simulations were executed using the DESMOND software package. Trajectory analysis and visualization were performed using XMGRACE and Discovery Studio Visualizer, respectively.

#### 1.5. Assessment of CD28/CD80 Interaction Disruption via Competitive ELISA

A competitive ELISA was performed using a CD28:B7-1[Biotinylated] Inhibitor Screening Assay Kit (Cat# 72007; BPS Bioscience, San Diego, CA, USA) to assess the inhibition of CD80 (B7-1) binding to immobilized CD28; manufacturer's instructions were followed and are briefly described here.. 96-well plates were coated with CD28 (2 µg/mL in PBS, 50 µL/well) and incubated overnight at 4°C. The following day, wells were washed with 1×Immuno Buffer and blocked with Blocking Buffer for 1 hour at room temperature. Test compound was added at varying concentrations along with B7-1-biotin (5 ng/µL) and incubated for 1 hour to allow competitive binding. Uncoated wells served as ligand controls, and inhibitor buffer was used in place of compound for control wells. After incubation, wells were washed, and Streptavidin-HRP (1:1000 in Blocking Buffer) was added for 1 hour. Chemiluminescent substrate was added after a final wash, and luminescence was measured immediately using a plate reader set to luminescence mode.

### 2. Chromatographic and Mass Spectra Data for Selected Small Molecule from the Discovery Diversity Set (DDS) Library

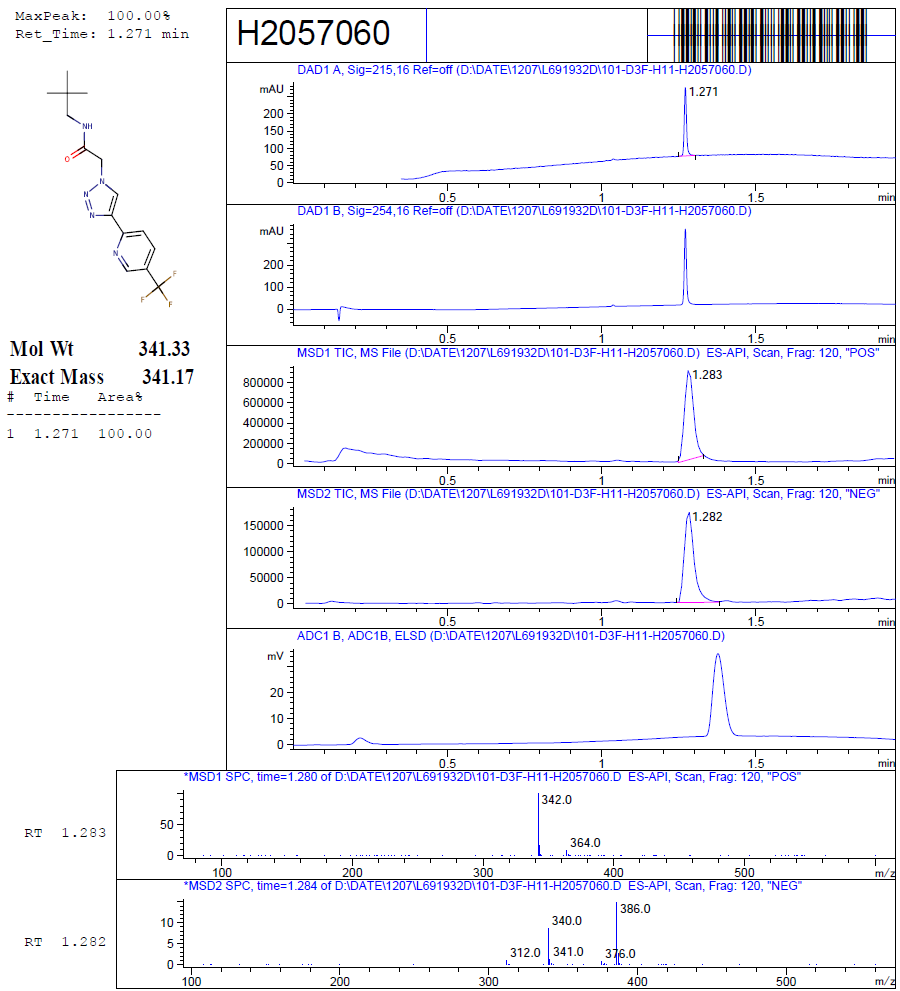

**Figure S1.** Chromatographic data and mass spectrum of **DDS1**.

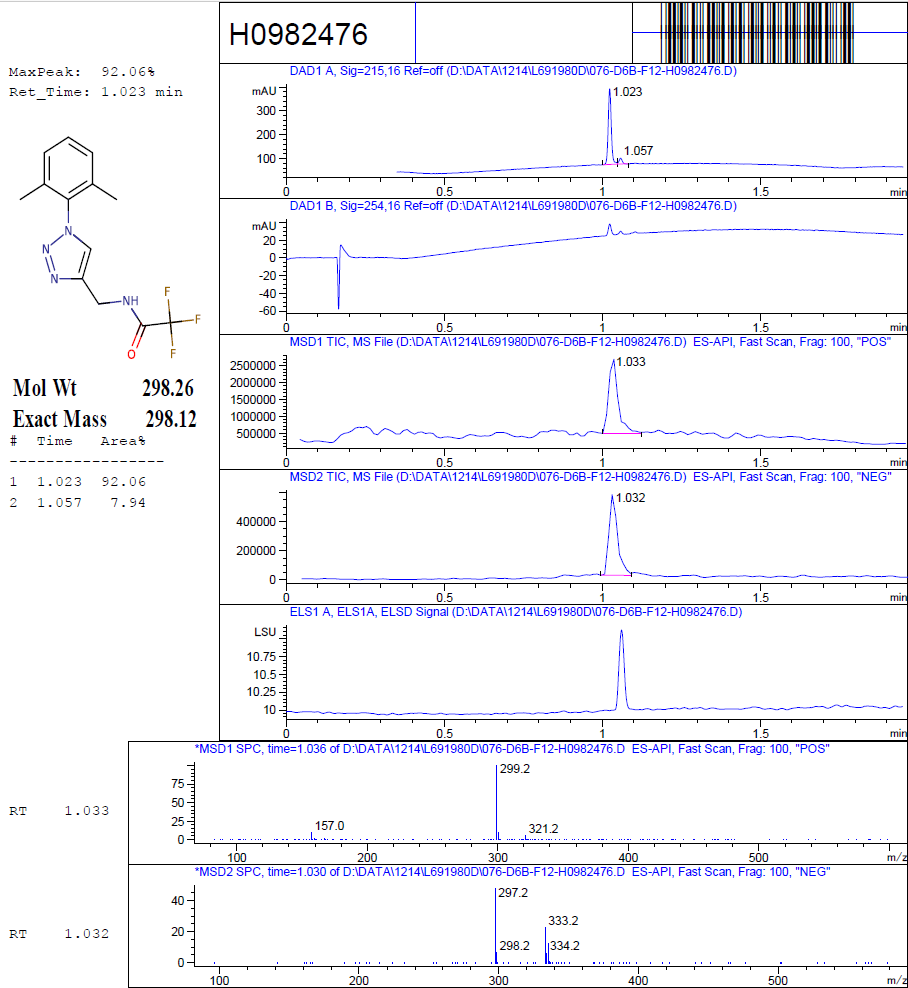

**Figure S2.** Chromatographic data and mass spectrum of **DDS2**.

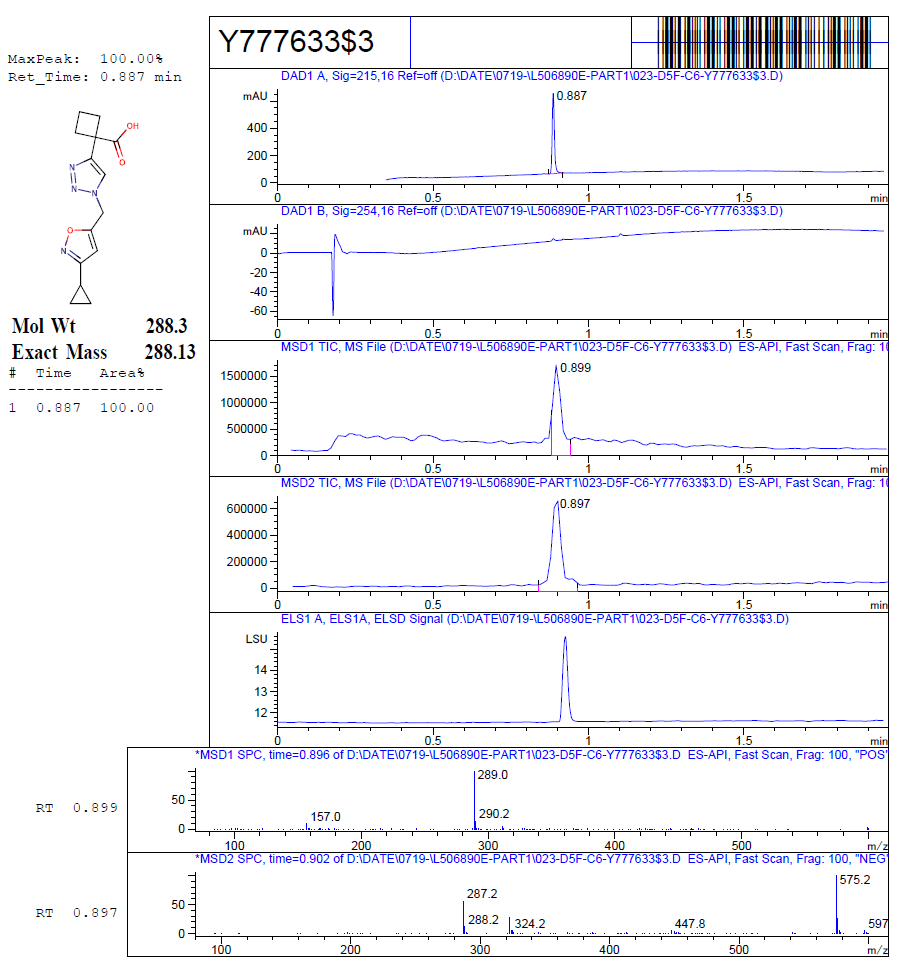

**Figure S3.** Chromatographic data and mass spectrum of **DDS3**.

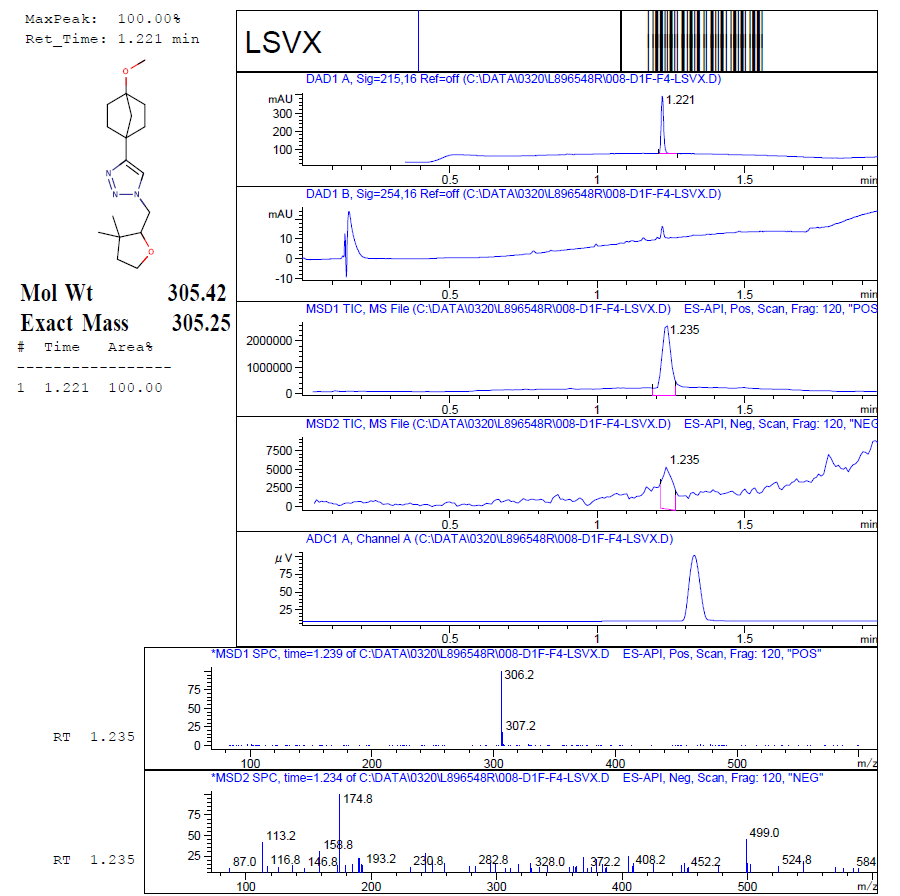

**Figure S4.** Chromatographic data and mass spectrum of **DDS5**.

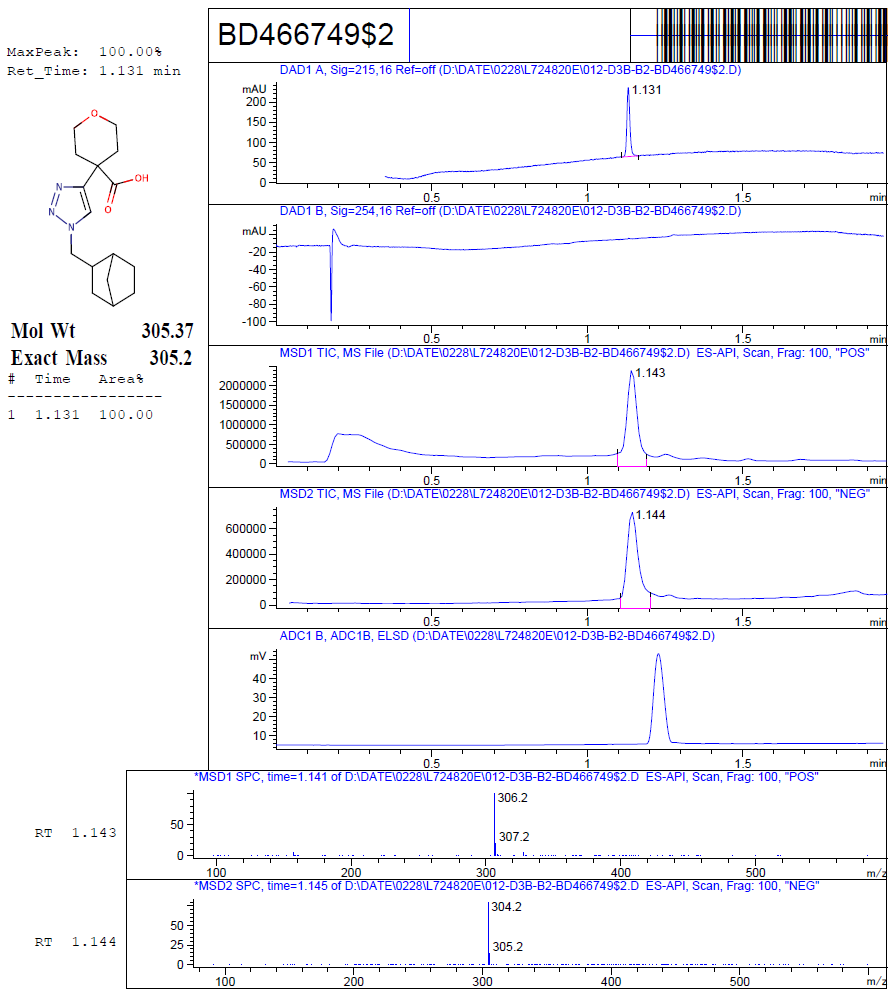

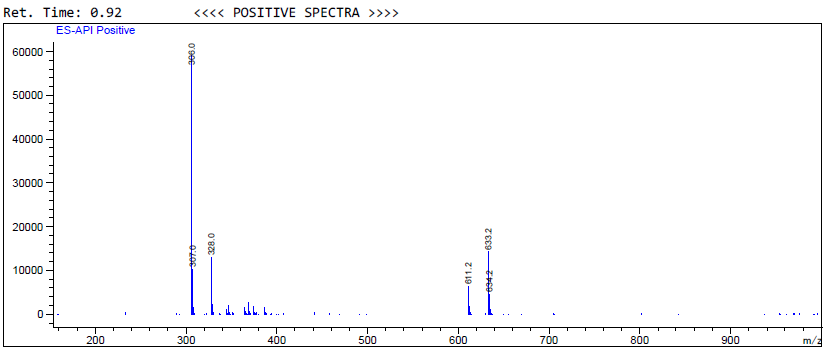

**Figure S5.** Chromatographic data and mass spectrum of **DDS6**.

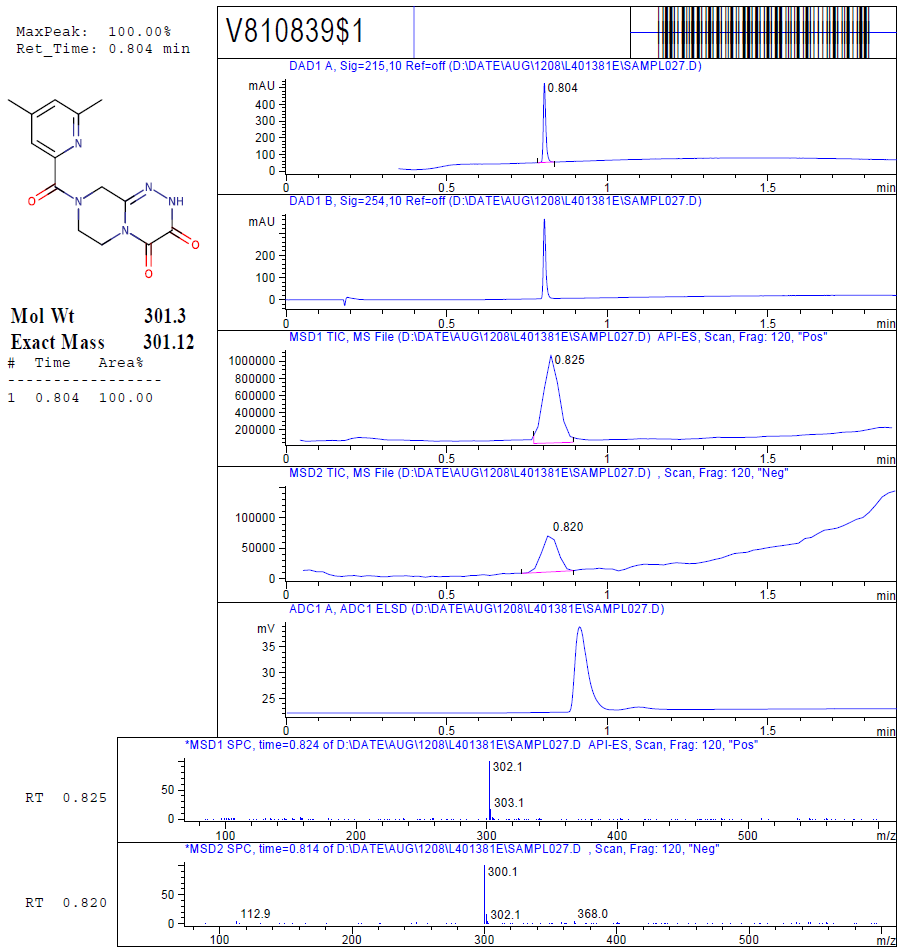

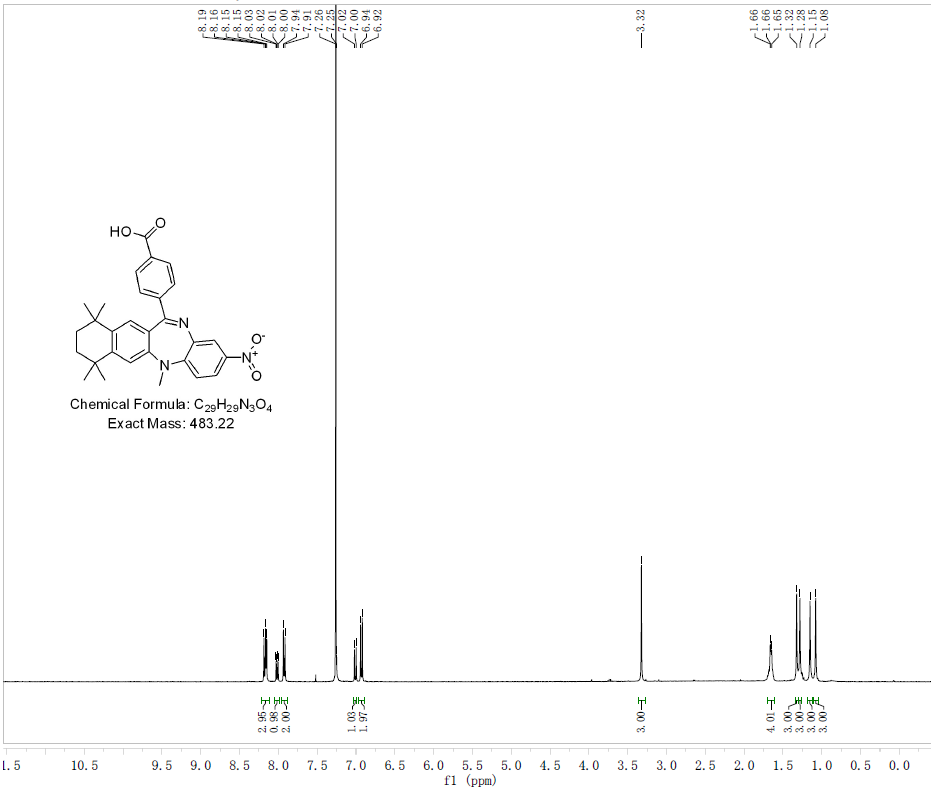

**Figure S6.** Chromatographic data and mass spectrum of **DDS7**.

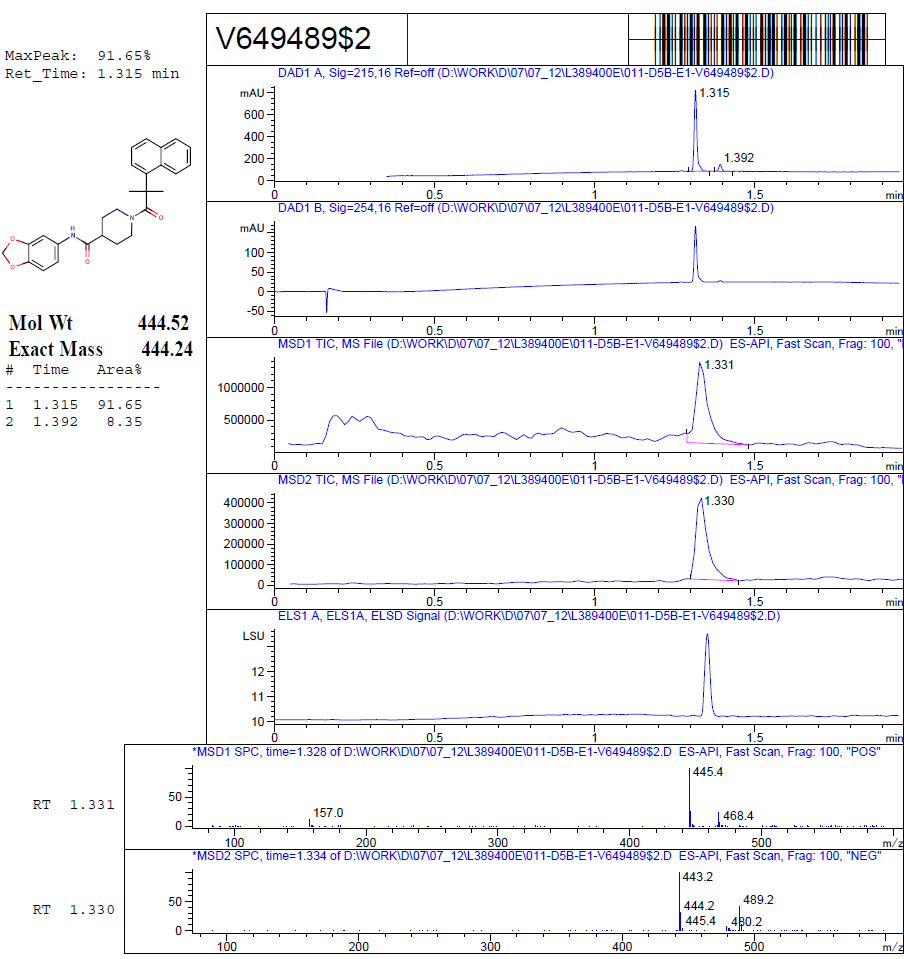

**Figure S7.** Chromatographic data and mass spectrum of **DDS8**.

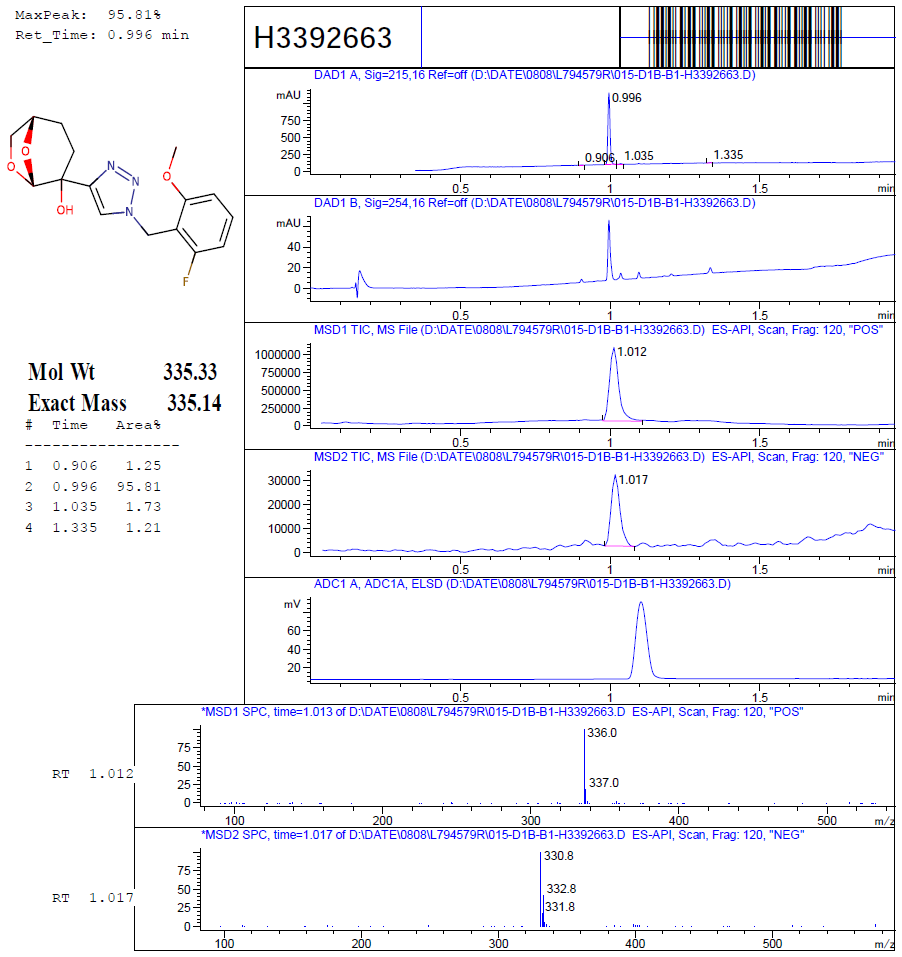

**Figure S8.** Chromatographic data and mass spectrum of **DDS9**.

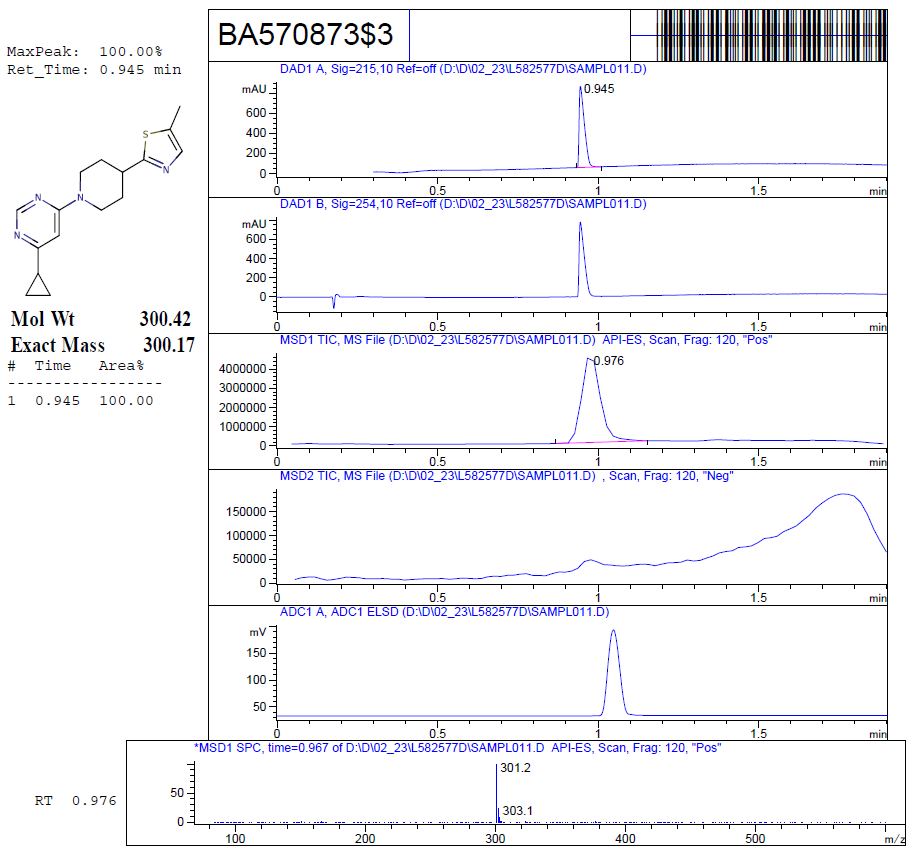

**Figure S9.** Chromatographic data and mass spectrum of **DDS10**.

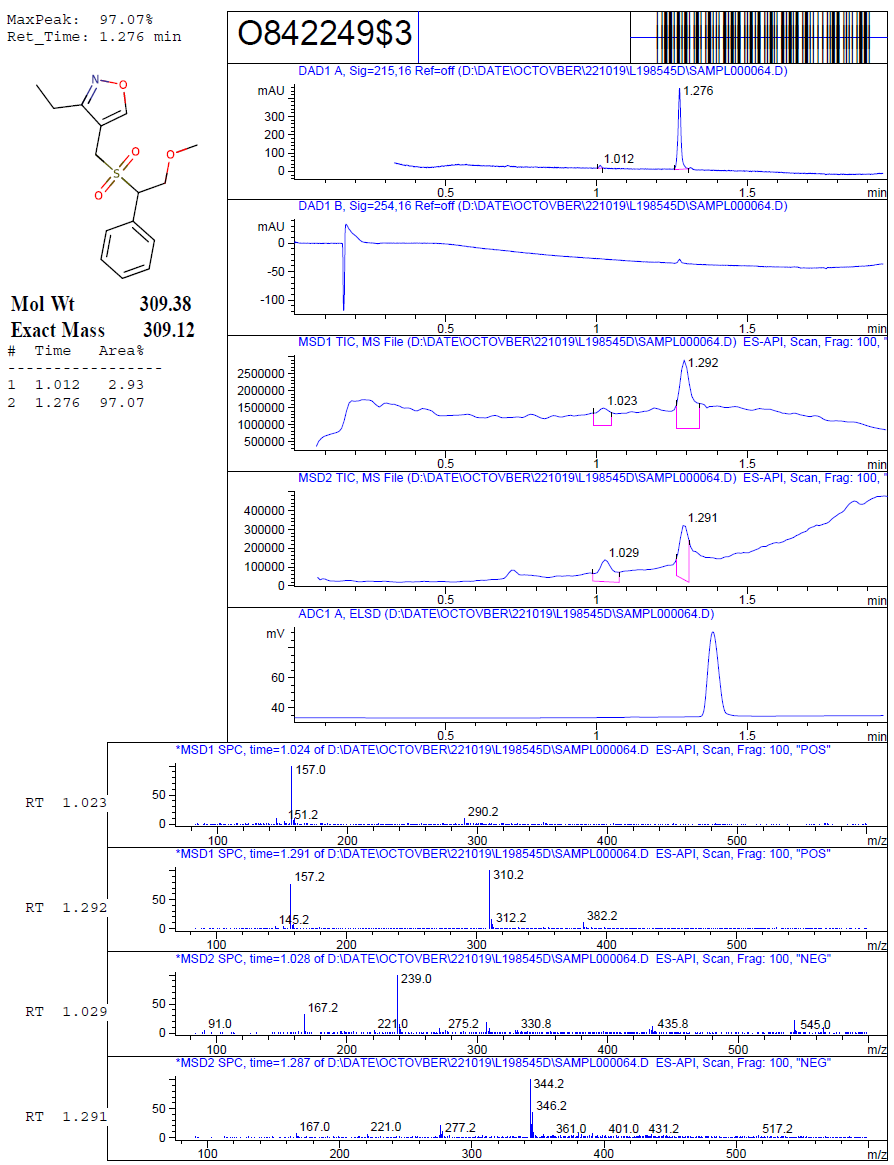

**Figure S10.** Chromatographic data and mass spectrum of **DDS11**.

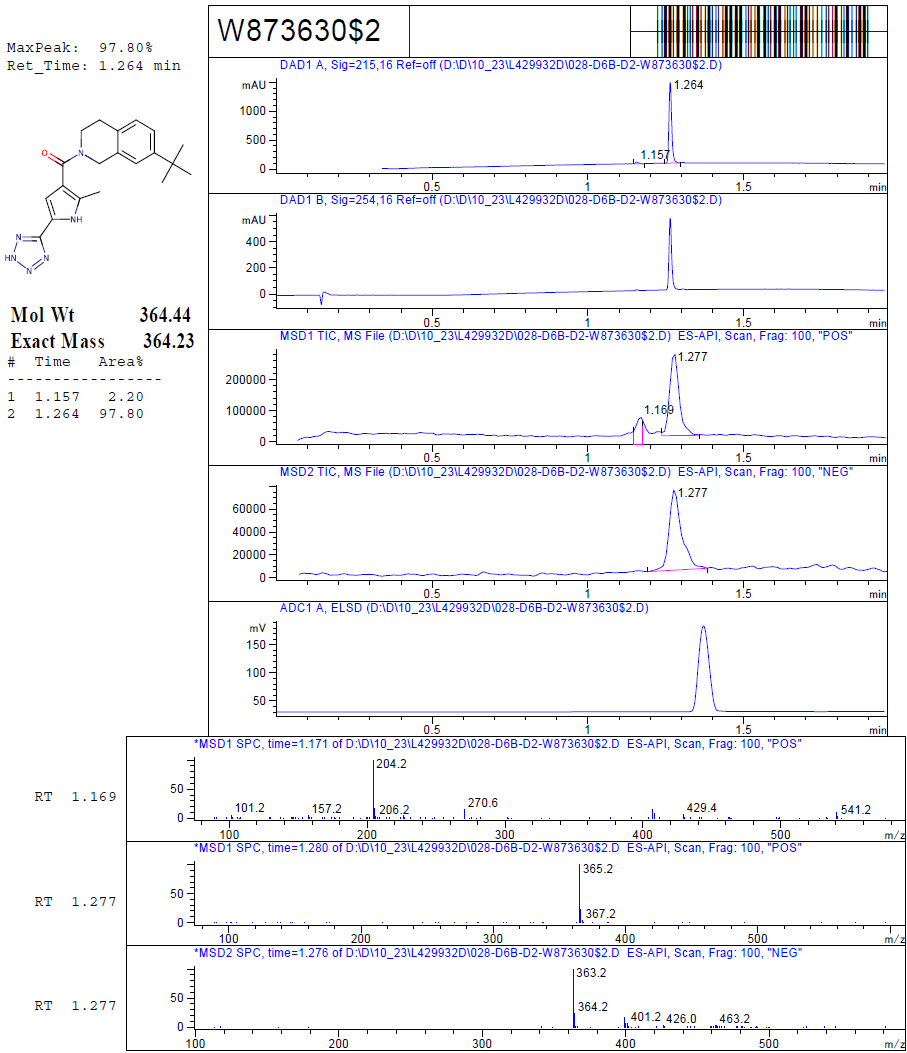

**Figure S11.** Chromatographic data and mass spectrum of **DDS12**.
